# DNA methylation supports accelerated biological age in Type 2 Diabetes which can be reversed with pharmacological treatments: Retrospective Cohort Study

**DOI:** 10.1101/2022.09.14.507908

**Authors:** Briana N. Cortez, Hui Pan, Cristina Aguayo-Mazzucato

## Abstract

**Background:** Biological age (BA) closely depicts age-related changes at a cellular level. Type 2 *Diabetes mellitus* (T2D) accelerates BA when calculated using clinical biomarkers. However, there is a large spread of individual BA within these groups and it is unclear what clinical biomarkers correlate with different speeds of aging and whether pharmacological treatment of diabetes alter BA. We hypothesized that accelerated BA would be seen at the DNA methylation (DNAm) level, the gold standard to determine BA, and biomarkers and treatments would correlate the rate of BA in T2D.

**Methods:** Publicly available DNAm samples were obtained from the GEO NCBI database and the NHANES 2017-2018 and ACCORD Cohorts were used for our analysis. We used the DNA Methylation Phenotypic Age algorithm and the Klemera and Doubal (KDM) methods to calculate BA with DNA methylation and clinical biomarkers, respectively.

**Results:** DNAm showed increased BA in whole blood and pancreatic islets in T2D in aging-related pathways, such as DNA damage and inflammation. Using the NHANES and ACCORD Trial cohorts, we found that avoidance of fried and fatty foods, and vigorous activity correlated with decreased BA in T2D. Cardiovascular, glycemic, and inflammatory biomarkers associated with the rate of aging in DM. Intensive blood pressure and T2D treatment associated with a greater deceleration in the speed of aging as compared to the standard.

**Conclusions:** Overall, we show that certain tissues age faster in people with T2D and this strongly associates with blood glucose control, inflammation and cardiovascular health. Effective treatment of the disease can decelerate aging and decrease BA suggesting the latter as a novel and integrated index to evaluate and follow people with T2D.

**Funding:** This study was supported by Institutional Startup Funds to C.A.M. (Joslin Diabetes Center) and NIH grants P30 DK036836 Joslin Diabetes Research Center (Bioinformatic Core).

## Introduction

Individuals with the same chronological age (CA), the time lived since birth, display great differences in their susceptibilities to age-related disease and death. The incidence of age-related disease continues to rise as the lifespan increases globally and great effort is being placed on understanding this phenomenon and increasing healthspan. The seven pillars of aging have been described as adaptation to stress, inflammation, epigenetics, macromolecular damage, metabolism, proteostasis, and stem cell population and regeneration^1^. The concept of biological age (BA) has emerged as a more precise measure of the cellular decline and physiological breakdown of the organs in the human body, and has been better correlated with an individual’s mortality and morbidity than CA^2^. For this reason, BA has great potential to be used in the clinic as a unique measure of a person’s risk for aging-related diseases and mortality. For example, biological age calculations based on specific organ systems, were found to have a higher correlation to disease severity than single biomarkers widely used in the clinic^3^. Many methods to calculate BA have been developed, including use of telomere length and clinical biomarkers. While use of readily available clinical biomarkers allows for greater efficiency in terms of cost and time, DNA methylation (DNAm) continues to be seen as the gold standard to calculate BA.

While many have studied BA in the individual irrespective of disease status, we showed that BA was significantly increased in *Diabetes melliuts (DM)* when using clinical biomarkers and the Klemera and Doubal method 1 (KDM)^4^. Individuals with Type 1 diabetes (T1D) were 16 years greater than those without diabetes and those with Type 2 diabetic (T2D) were 12 years than age and gender matched controls. Additionally, we found that a 9.85 year increase in BA correlated with a significant increase in mortality^4^. While an increase in BA in those with DM at the population level was shown, there was a large spread of individual BA, suggesting that additional variables were contributing to accelerated aging in DM.

We questioned whether calculating BA based on DNAm using the Phenotypic Age (DNAm PhenoAge) clock would corroborate our results showing increased BA in T2D. Additionally, we questioned whether specific biomarkers, lifestyle factors, individual characteristics, and treatment regimens were associated with the rate of BA using the Action to Control Cardiovascular Risk in Diabetes (ACCORD) and National Health and Nutrition Examination Survey (NHANES) Cohorts.

Overall, we found that T2D individuals have accelerated aging in pancreatic islets and whole blood but not in adipose or skeletal muscle using DNAm PhenoAge. Associations between lifestyle, individual characteristics, cardiovascular, glycemic, and lipid biomarkers, had a significant association with the rate of BA in DM. Pharmacological treatment of hyperglycemia, blood pressure and hyperlipidemia after a 48 month period, returned accelerated BA to the same or lower levels than CA suggesting that BA is modifiable as well as a novel and integrated index to evaluate and follow people with T2D and their treatment.

## Methods

### Study Population

This study was approved by the Joslin Diabetes Center’s Committee on Human Studies (CHS) which determined that it represents exempt human subject research under 45 CFR 46.104 (d)(4)(ii): it involves the secondary research use of previously collected identifiable private information which was recorded in such a manner that the identity of the human subjects cannot be ascertained directly or through identifiers.

Deidentified patient data was obtained from the Action to Control Cardiovascular Risk in Diabetes (ACCORD) Trial and the National Health and Nutrition Examination Survey (NHANES) 2017–2018 Cohort. Participation in the ACCORD trial required the diagnosis of T2D in individuals greater than 40 years old, who may have also had coexisting hypertension (HTN) and/or hyperlipidemia (HLD). All participants of the ACCORD trial were randomized into a standard or intensive glycemia cohort. Those who had HTN were also randomized to a standard or intensive blood pressure (BP) trial, and those with HLD were randomized into the lipid fibrate or placebo trial. Individuals in the lipid fibrate group received a statin and fibrate, and those in the lipid placebo group were treated with a statin and placebo. The target hemoglobin A1c (HbA1c) for those in the intensive glycemia group was less than 6%and 7.0-7.9%, for the standard group. The target systolic blood pressure (SBP) for the intensive group was below 120 mmHg, and below 140 mmHg for the standard group. Individuals were placed on a variety of oral medication for glucose and blood pressure control in order to meet their target SBP and/or HbA1c values.^5^

We calculated BA using KDM from individuals enrolled in the ACCORD trial at baseline, 48 months, and at their exit point. The mean follow up time was 4.7 years.^5^ Those with absent biomarkers at any of the 3 times points were excluded.

Individuals from the NHANES Cohort were 20-70 years old and grouped as nondiabetic, prediabetic or diabetic. Those who stated that they did not have diabetes, were placed in the nondiabetic population. Because the American Diabetes Association guidelines classify prediabetes as a hemoglobin A1c (HbA1c) of 5.7 to 6.4%, those with values in this range were excluded from the nondiabetic group to eliminate participants with undiagnosed prediabetes or DM.^6^

### Calculation of Biological Age

#### DNA Methylation Phenotypic Age

The GEO NCBI database provided access to DNA methylation data from people with and without T2D. We accessed samples from pancreatic islets (GSE21232), whole blood (GSE34008), subcutaneous adipose and skeletal muscle (GSE38291), processed using Illumina Infinium Human Methylation 27 Bead Chip.^7,8^. The distribution of tissue samples is shown in **Table 1, Source 1**. In all tissues, the age differences were not statistically significant between nondiabetic and T2D groups.

**Table 1, Source 1:** Tissues obtained from GEO NCBI Database used for DNA methylation analysis.

| Tissue | NCBI Geo Database Identifier | Nondiabetic Samples | T2D Samples |
| --- | --- | --- | --- |
| Pancreatic Islets | GSE21232 | 11 | 5 |
| Whole Blood | GSE34008 | 12 | 12 |
| Muscle | GSE38291 | 11 | 11 |
| Adipose | GSE38291 | 5 | 5 |

DNA Methylation Phenotypic Age (DNAm PhenoAge) was used to calculate BA due to its higher reported accuracy.^9^ DNAm PhenoAge was run in R using the methyl package at https://github.com/isglobal-brge/methylclock.^10^ After calculation, CpG sites that were statistically different between nondiabetic and T2D samples were matched to gene symbols, and Pathway analysis was completed using Reactome.

### Klemera and Doubal Method

Selection of biomarkers and calculation using the Klemera and Doubal Method 1 (KDM) within this study followed the same methodology defined in our previous study^4,11^. A total of 7 biomarkers were used in this algorithm including age (years), serum creatinine (mg/dL), albumin (g/dL), cholesterol (mg/dL), systolic and diastolic blood pressure (mmHg), pulse (beats per minute), and glycohemoglobin (%). Due to the absence of serum albumin in the ACCORD Cohort, this variable was excluded from the KDM calculation and only the 6 remaining variables were included. Winsorization was performed in analysis of the NHANES cohort. Results from the ACCORD study were expressed in terms of age ratio, with the exception of diet, race, and biological sex analysis. This algorithm was processed in R studio, using the bioage package at https://github.com/bjb40/bioage.

### Definition of Age Ratio and dAge

Age ratio is calculated as the ratio between BA and CA. An age ratio equal to 1, indicates that both BA and CA equal. If age ratio is greater than 1, BA is greater than CA and vice versa when less than 1.

Delta age (dAge) is calculated as the difference between BA and CA. When equal to 0, BA and CA are equal. A positive value indicates that BA is greater than CA, and vice versa when the value is negative.

### Definition of fast, slow, and normal aging individuals

Individuals were classified in three groups based on the correlation between their BA and CA: 1) Fast aging individuals, or fast agers, are those with a BA greater than 5 years from their CA; 2) Slow aging individuals, or slow agers, are those with a BA less than negative 5 years from their CA; 3) Normal agers are those whose BA is within 5 years plus or minus their CA.

### Definition of obesity with use of dual-energy x-ray absorptiometry (DEXA)

Body mass index (BMI) is the measure currently used to determine normal, overweight and obesity status in patients. DEXA scan technology may be used to determine fat percentage. Currently, there is no standard range used to subclassify individuals based on obesity status with use of DEXA. We classified those with total body fat percentage of less than 25, as normal weight individuals. Those with a total body fat percentage between 25-30 were considered overweight, and those with a percentage greater than 30 were considered obese.

### Statistical Analysis

Multiple linear regression and nonparametric t-test was performed using R, Microsoft Excel and Prism in order to find associations between biomarkers and treatment groups and rate of BA. If applicable, normal clinic ranges were indicated on graphs by gray shaded areas. The number of patients in each group are indicated in the figure or figure legends.

## Results

### DNA Methylation Phenotypic Age correlated with increased BA

To further understand our previous results showing an increased BA in those with T2D,^4^ DNAm PhenoAge was used to calculate BA in people with T2D. Publicly available DNA methylation samples from the GEO NCBI Database were compiled from whole blood^8^, pancreatic islets^8^, adipose^7^ and skeletal muscle^7^.

When comparing nondiabetic and T2D samples, we found that DNAm phenotypic age from people with T2D was increased in whole blood by 10 years, and pancreatic islets by 15 years. However, there was no difference between T2D and nondiabetic samples in adipose and skeletal muscle (**Figure 1A-D, Source 1**).

**Figure 1, Source 1:**
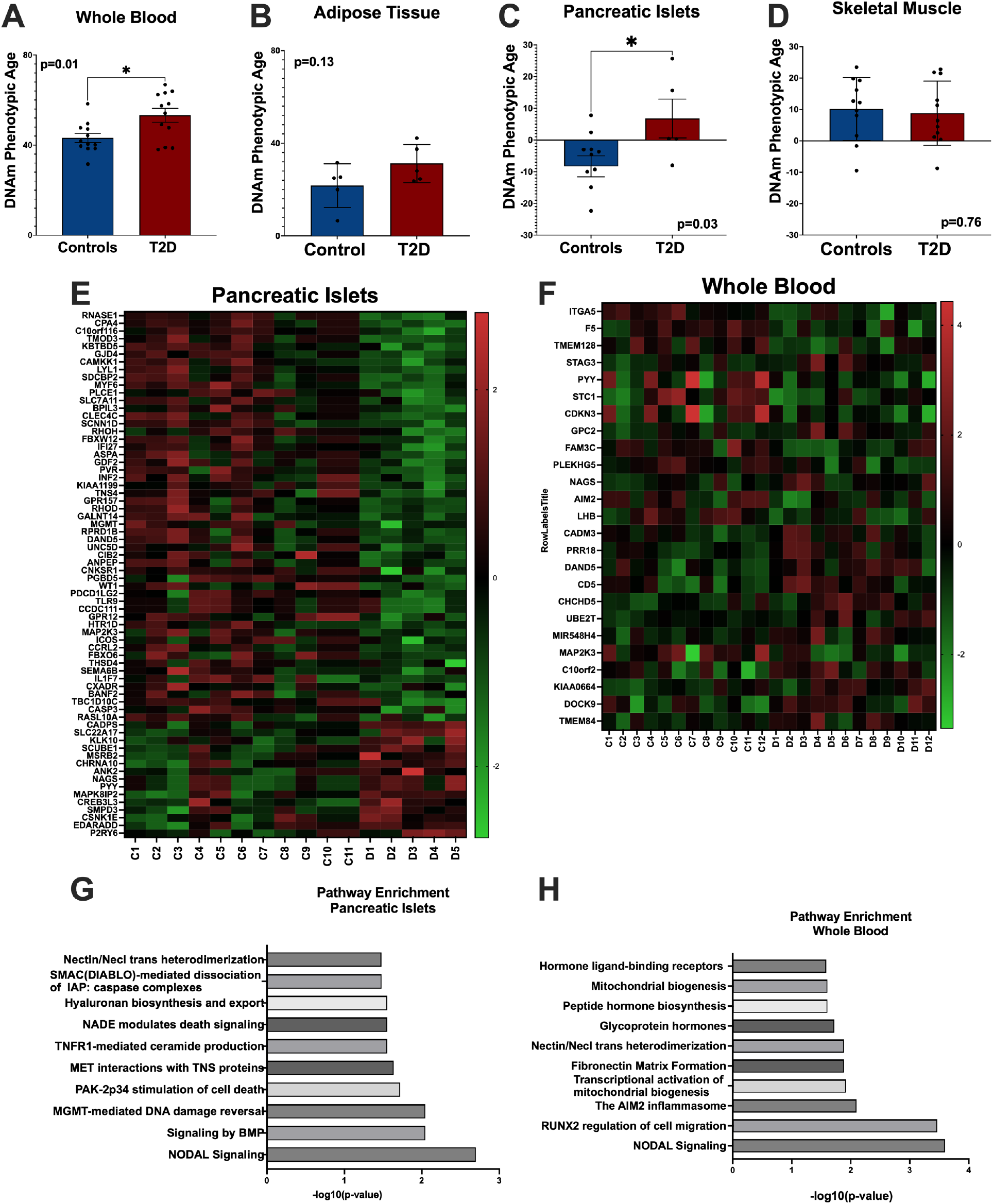
DNA methylation phenotypic age correlates with increased phenotypic age in T2D in whole blood (A) and pancreatic islets (B). No correlation in is shown adipose tissue (C) and muscle tissue (D). Nonparametric t-test was performed. Heat maps show differences in gene regulation in pancreatic islets (E), and whole blood (F). Pathway enrichment of pancreatic islets (G), and whole blood (H) show similarities and differences in terms of pathway upregulation. *p<0.05, ** p<0.01, *** p<0.001

Pathway analysis of the CpG sites that were differentially methylated in pancreatic islets and whole blood samples showed genes associated with the hallmarks of aging.^1^ Pancreatic islet genes were significantly associated in pathways including signal transduction (*SDCBP2, PLCE1, RHOH, GDF2, KIAA1199, GPR157, CNKSR1*), DNA repair (*MGMT, RPRD1B, CCDC111, CSNK1E*), and programed cell death (*PDCD1LG2, CASP3, EDARADD*). Whole blood was significantly associated with genes involved in immune system function (*PLEKHG5, CD5*), metabolism of proteins (*NAGS, UBE2T*), organelle biogenesis and maintenance (*CHCHD5, KIAA0664, GPC2*) (**Figure 1G, H, Source 1**).

### Specific biomarkers correlated with speeds of aging

Given the individual variation of BA in people with T2D, the NHANES diabetic cohort was grouped into fast, slow, and normal agers, and analysis of biomarkers with multiple linear regression was completed. Both, cardiovascular parameters and glycemic indices were correlated with speeds of aging (**Figure 2, Source 2**). Increased levels of systolic blood pressure (**Figure 2A, Source 2**), diastolic blood pressure (**Figure 2A, Source 2**), pulse (**Figure 2B, Source 2**), and decreased albumin (**Figure 2C, Source 2**) correlated with accelerated aging in people with T2D. We used albumin as a cardiovascular parameter due to its role in the regulation of the viscosity of blood and fluid dynamics. Increased levels of glycohemoglobin (**Figure 2D, Source 2**) and fasting glucose (**Figure 2E, Source 2**) correlated with faster aging. Those who were first diagnosed with diabetes at an older age, were correlated with slower speeds of aging (**Figure 2F, Source 2**).

**Figure 2, Source 2:**
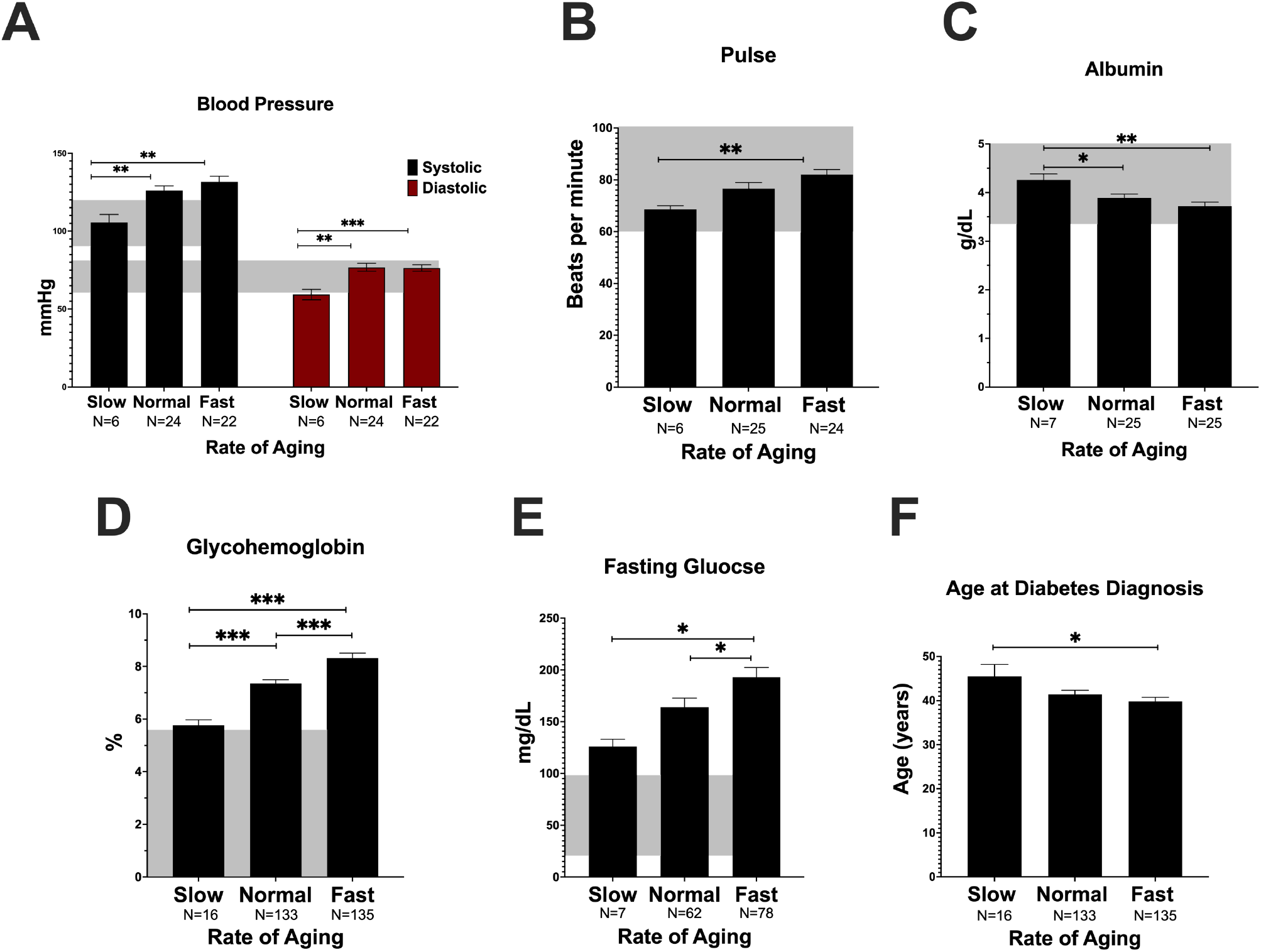
Cardiovascular and glycemic measures correlate with slow (n=16), normal (n=133) and fast (n=135) agers including systolic and diastolic blood pressure (A) pulse (B) albumin (C) glycohemoglobin (D), fasting glucose (E). Older age at diabetes diagnosis correlates with faster aging (F). Normal clinic ranges were indicated on graphs by shaded areas. Nonparametric t-test was used for statistical analysis. * p<0.05, ** p<0.01, *** p<0.001. Shaded areas represent normal clinical ranges.

Analysis of lipid and adiposity parameters in the NHANES diabetic cohort also correlated with alterations in speed of aging (**Figure 3, Source 3**). Increases in total (**Figure 3A, Source 3**) and LDL (**Figure 3B, Source 3**) cholesterol were correlated with increased speeds of aging. Normal agers had an HDL level that was significantly lower than fast agers (**Figure 3B, Source 3**). While obesity status was not correlated with BA in nondiabetic and prediabetic populations, it was correlated with increased dAge in the diabetic population (**Supplemental Figure 3, Figure 3C, Source 3**). Use of DEXA scans showed a significant increase in total fat percentage in diabetic individuals compared with nondiabetic individuals (**Figure 3D, Source 3**). Total trunk fat was significantly increased in diabetic and prediabetic individuals compared to nondiabetic individuals (**Figure 3E, Source 3**). Bilateral upper extremities of individuals with diabetes had a significant increase in fat percentage (**Figure 3G, Source 3**) as compared to nondiabetic individuals. Bilateral lower extremities were not associated with differences in fat percentages between nondiabetic, prediabetic or diabetic individuals (**Figure 3F, Source 3**).

**Figure 3, Source 3:**
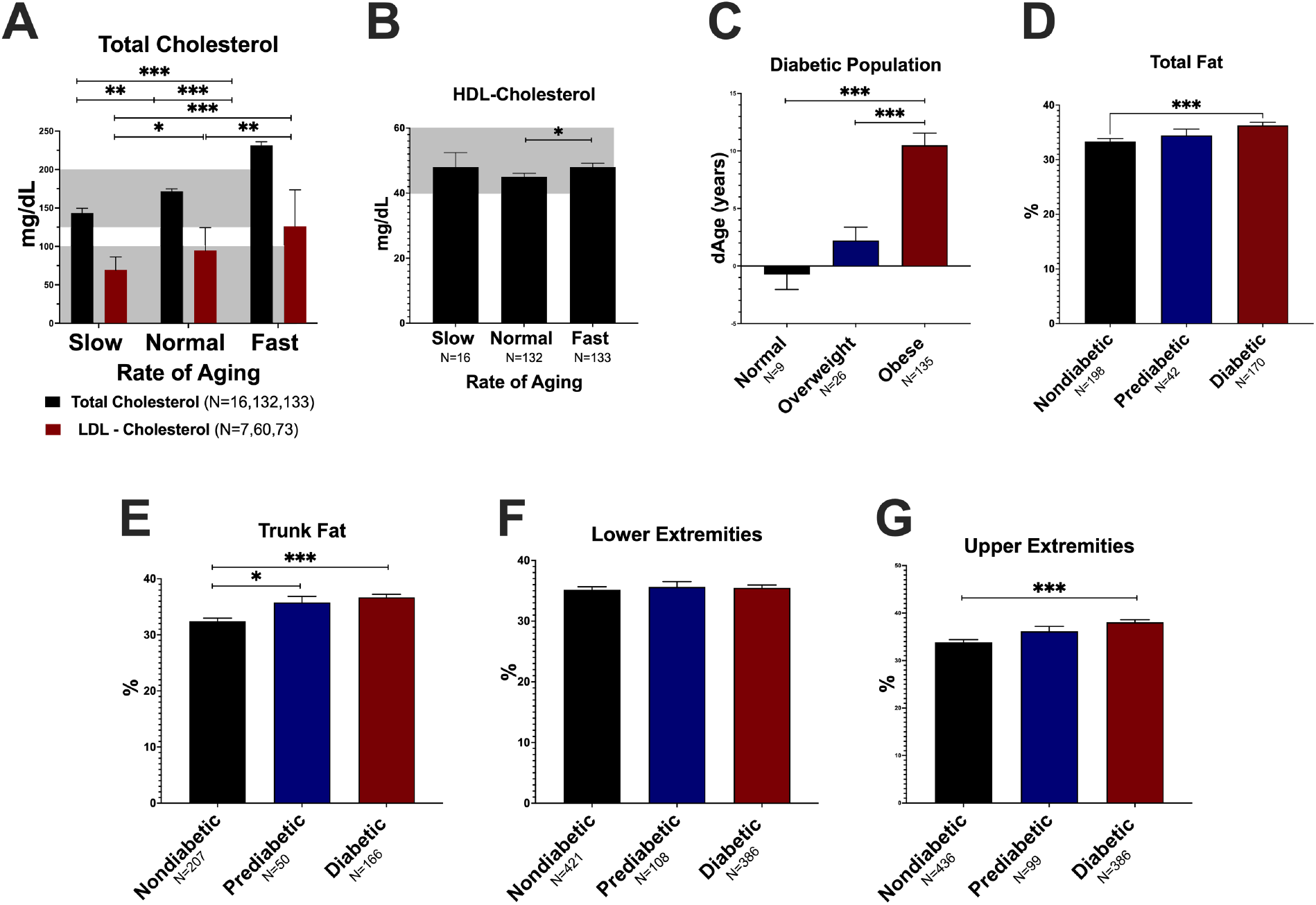
Lipid markers correlate with slow, normal, and fast agers increased aging in total and LDL cholesterol (A), and HDL cholesterol (B). The diabetic population was correlated with increased dAge in overweight and obese individuals (C). Increased percentage of total fat (D), trunk fat (E), and upper extremity adiposity (G) correlate with diabetes, while lower extremity adiposity (F) does not. *p<0.05, ** p<0.01, *** p<0.001. Shaded areas represent normal

Given the importance of inflammation in aging^12^, whole blood and inflammatory markers were analyzed for associations with speeds of aging (**Figure 4, Source 4**). Increased globulin levels and lower mean cell (Hb) concentration correlated with increased speeds of aging (**Figure 4A, Supplemental Figure 2, Source 4, 1s**). Normal aging individuals had a mean ferritin level of 213 ng/dL, and lower levels correlated with faster aging (**Supplemental Figure 2, Source 1s**). Increased number of platelet counts, and segmented neutrophils were correlated with faster speeds of aging (**Figure 4B, C, Source 4**), both of which are indicators of systemic inflammation.

**Figure 4, Source 4:**
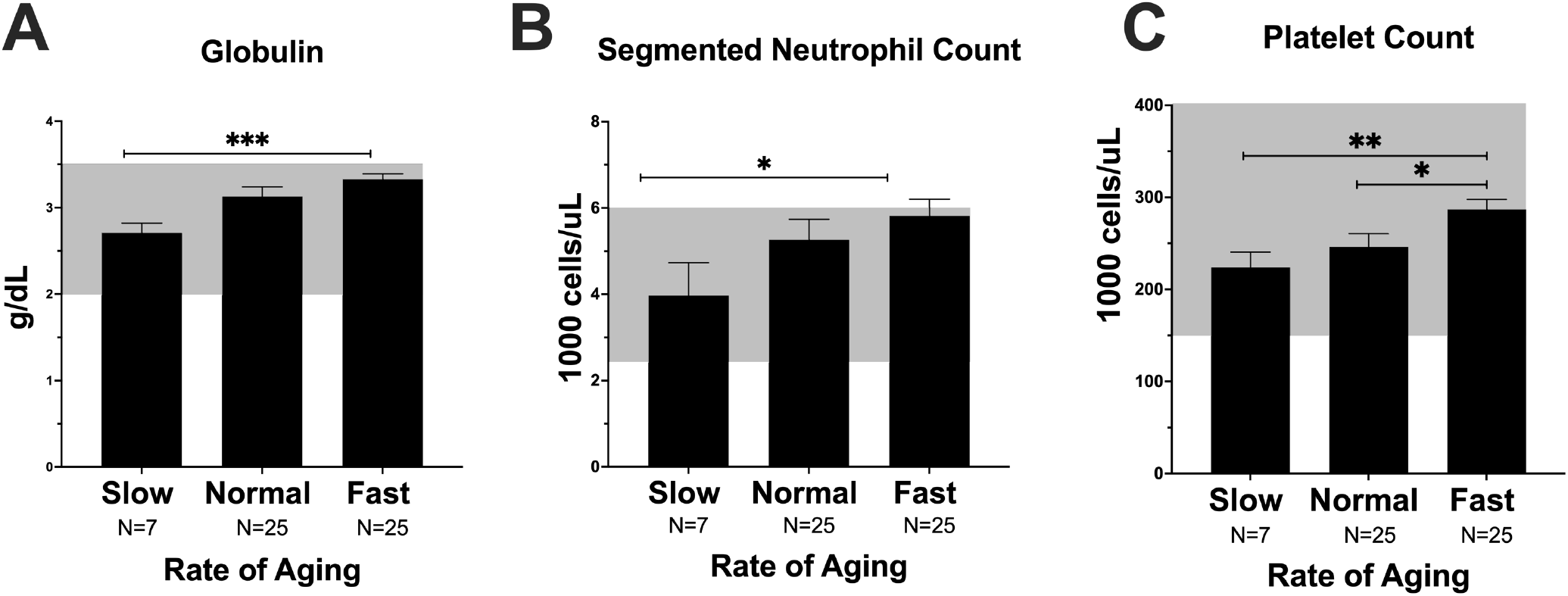
Inflammatory markers correlate with increased aging, including globulin (A), segmented neutrophil count (B), and platelet count (C). *p<0.05, ** p<0.01, *** p<0.001. Shaded areas represent normal clinical

### Lifestyle and population characteristics correlated with speed of aging

Lifestyle factors were evaluated for correlations with speeds of aging. In the NHANES cohort lower dAge in nondiabetic individuals was correlated with walking and biking (**Figure 5B, Source 5**), while vigorous and moderate intensity activities were not (**Supplemental Figure 3A, B, Source 5**). In the diabetic population however, only vigorous exercise was correlated with a lower dAge, whereas walking and biking, and moderate activity were not (**Supplemental Figure 3C, D, Source 5**). None of the intensities of exercise correlated with dAge in those with prediabetes (**Supplemental Figure 3E, F, G, Source 5**).

**Figure 5, Source 5:**
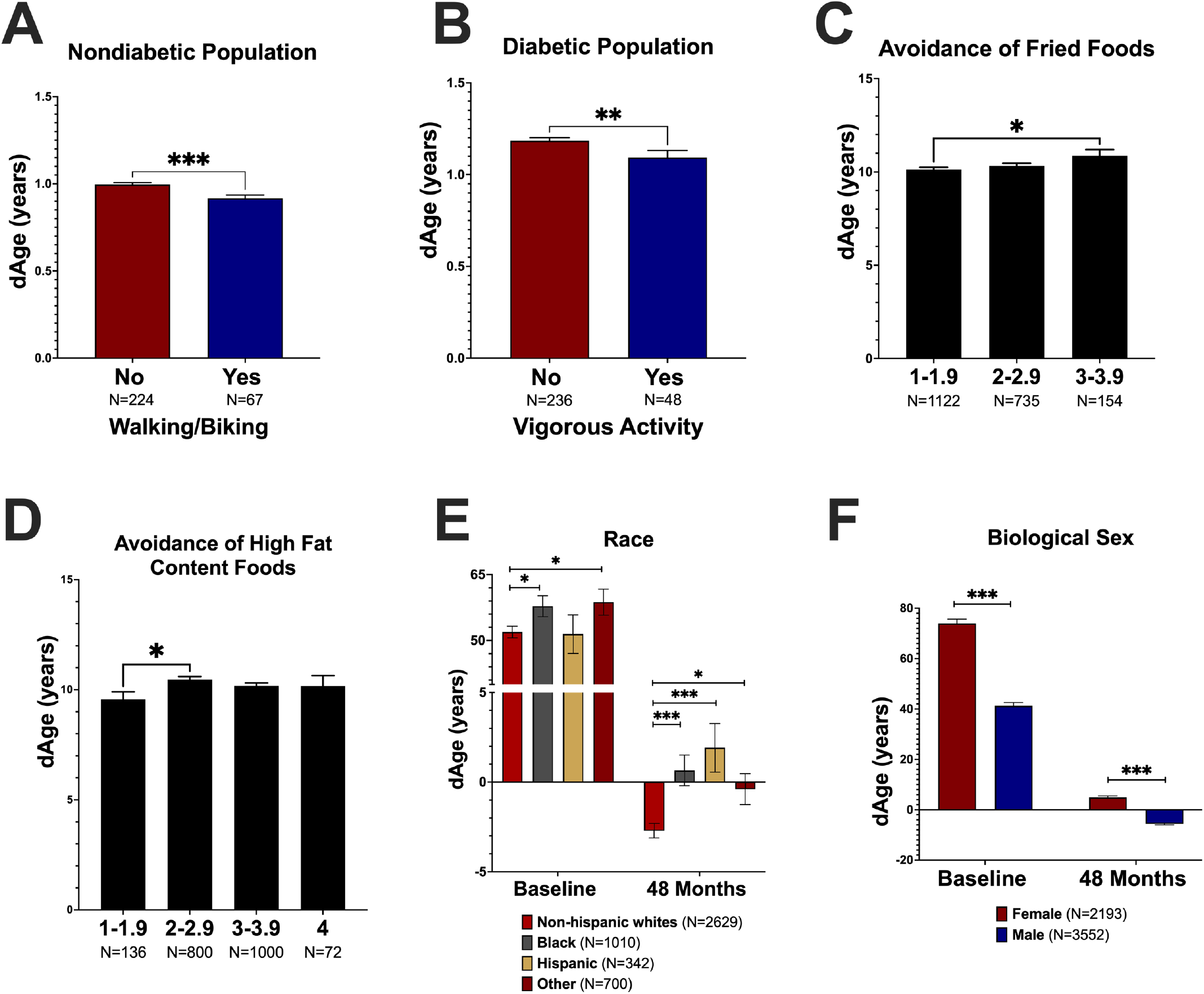
Lifestyle factors and population characteristics correlate with differences in rates of aging. Walking and biking in nondiabetics (a) and vigorous activity in diabetics (b) are correlated with decreased dAge. Avoidance of fried (c) and high fat content (d) foods correlate with lower dAge. A greater deceleration in rate of aging between races was only seen in whites vs. Hispanics (E). Both biological sex had a significant decrease in their dAge from baseline to 48 months of enrollment in the study, however males had a significantly lower dAge at baseline and 48 months when compared to females (F). *p<0.05, ** p<0.01, *** p<0.001

When assessing diet within the T2D ACCORD population, avoidance of fried (**Figure 5C, Source 5**) and high fat content foods (**Figure 5D, Source 5**) correlated with a lower dAge (a rating of 1 being most avoidance and 5 least avoidance). When asked to subjectively rate their overall health, fast agers in the NHANES study reported a poorer perception of their health (**Supplemental Figure 4, Source 1s**).

Comparisons between race and biological sex were performed using BA calculated at baseline and 48 month time points within the ACCORD Study. Overall, all cohorts of race and biological sex had significant decelerations in rate of aging over the 48 months (**Figure 5E, F, Source 5**). Blacks had a greater dAge, which indicates higher BA, at both baseline and 48 months when compared to whites. Hispanics had no significant difference at baseline but had a significantly higher dAge at 48 months as compared to whites. This indicates that the Hispanic cohort had a lower magnitude of deceleration in their rate of aging as compared to whites (**Figure 5E, Source 5**) and that Blacks had higher BA throughout the whole study. In terms of biological sex differences, females in the study had a higher BA/CA ration throughout the study indicating faster biological aging (**Figure 5F, Source 5**).

### Glycemic and blood pressure pharmacological control correlated with decreased aging

BA was calculated using KDM in individuals from the ACCORD cohort at baseline, 48 months, and at exit from the study and rates of aging were compared between cohorts. Within the standard glycemia cohort, those dually enrolled in the intensive BP trial had a significantly greater reduction in their rate of aging over 48 months as compared to the standard (**Figure 6A, Source 6**). Those who were enrolled in the standard glycemia cohort and dually enrolled in the lipid fibrate cohort, had a greater deceleration in BA when compared to the lipid placebo cohort (**Figure 6B, Source 6**).

**Figure 6, Source 6:**
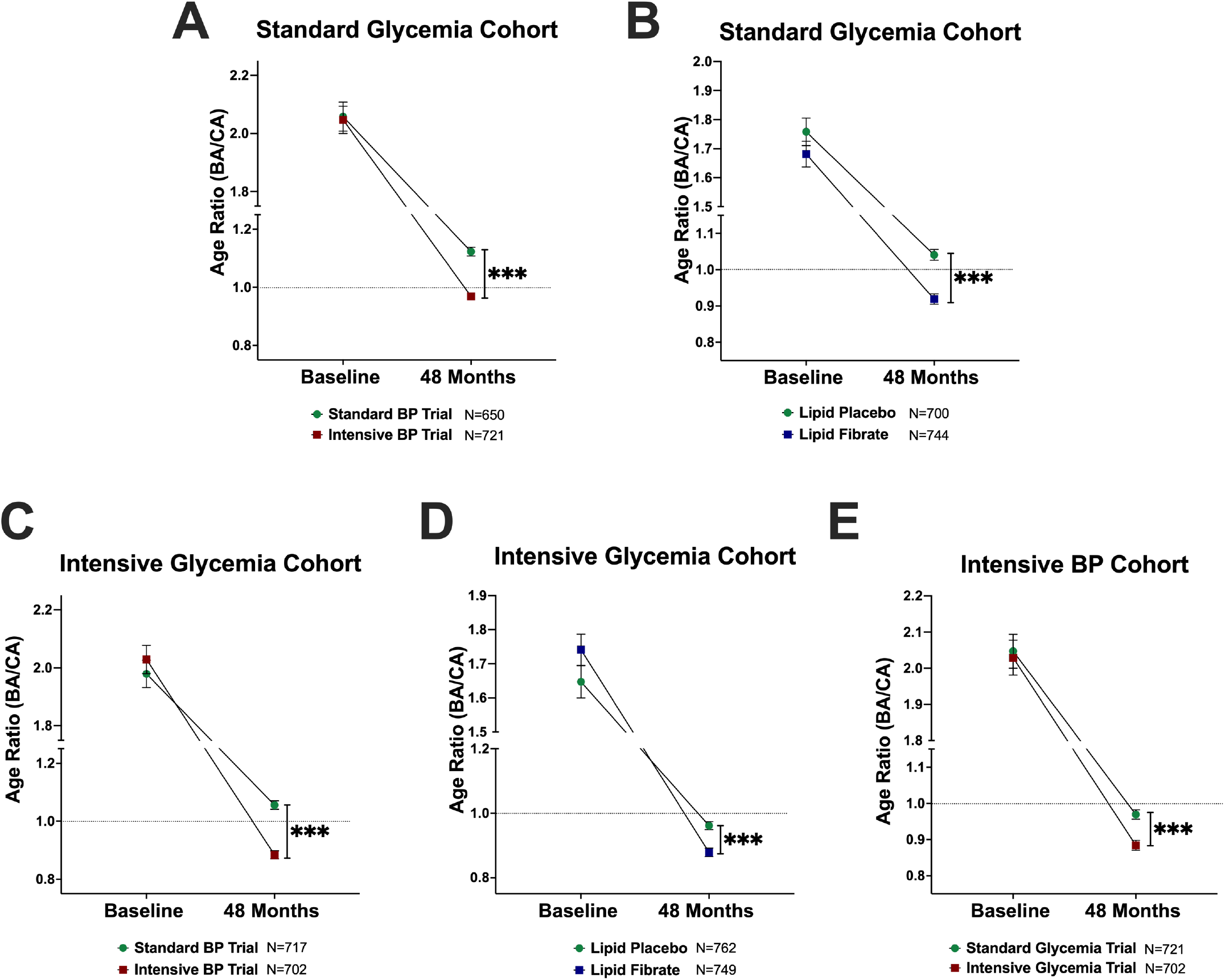
Rate of aging comparisons between ACCORD cohorts after 48 months in the standard glycemia (A and B), intensive glycemia (C and D), and standard blood pressure (E). Analyzed using nonparametric t-test. *p<0.05, ** p<0.01, *** p<0.001

Those in the intensive glycemia cohort who were dually enrolled in the intensive BP trial, or the lipid fibrate trials had a significantly greater reduction in their rate of aging over 48 months when compared with the standard and lipid placebo cohorts (**Figure 6C, Source 6**).

Those in the standard BP cohort who were also enrolled in the intensive glycemia trial had a greater deceleration in BA when compared to the standard glycemia trial (**Figure 6E, Source 6**). Further comparisons were made between standard and intensive BP and glycemia trials with lipid fibrate and placebo trials. The same trend remained; overall intensive treatments with lipid fibrate treatment were more successful in reducing BA over 48 months (**Supplemental Figure 4, Source 6**).

Overall, these results show that whereas at the beginning of the study the BA of all individuals were significantly greater than their CA, pharmacological treatment of diabetes, hypertension and hyperlipidemia was able to bring BA to CA levels or even below. More intensive treatment of all three clinical conditions, achieved a greater reduction in BA.

### Glycemia medications correlated with decelerated aging

We assessed whether specific medications to treat T2Dcorrelated with rates of aging. In the ACCORD Trial, insulin and 5 classes of glycemic oral medications were used including metformin, thiazolidinedione, sulfonureas, meglitinide, and a-glycosidase inhibitors. All oral medications and insulin decreased age ratio from baseline to exit, no medication was more beneficial than others at reducing age ratio (**Figure 7A, Source 7**). The differences in age ratio between medication groups was already present at baseline, raising the possibility that BA at presentation might influence indication for specific medications further underscoring the importance of its routine consideration in the evaluation and follow up of people with T2D.

**Figure 7, Source 6:**
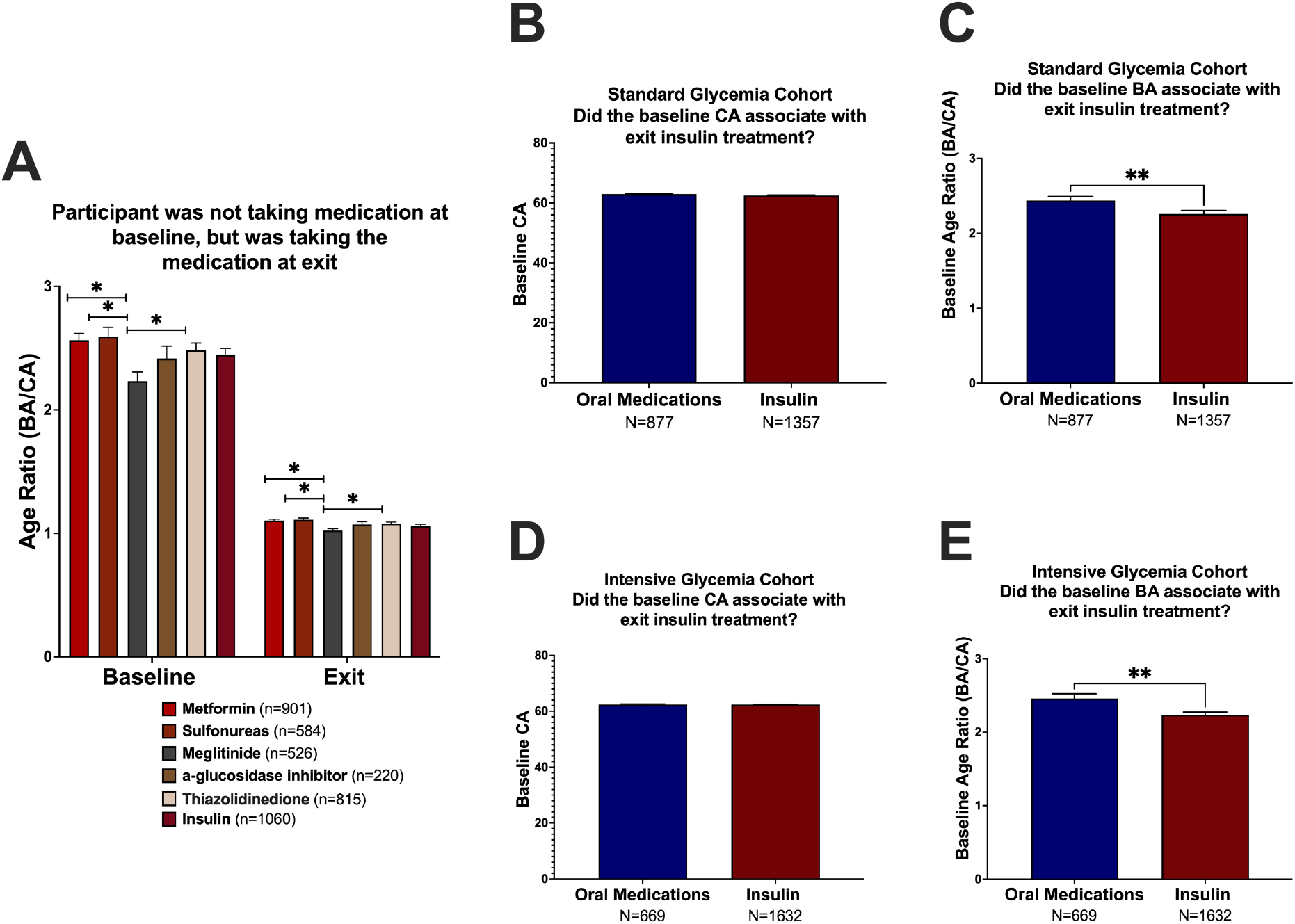
Significant decrease in age ratio from baseline to exit in patients taking any glycemic medications, however there was no significant difference between rates of aging from baseline to exit between medications when adjusted for significance at baseline (A). There was no difference in baseline CA in the standard glycemia (B) cohort, and higher baseline age ratio was correlated with lower use of insulin the exit time point (C). There was no difference in baseline CA in the intensive glycemia (D) cohort, and higher baseline age ratio was correlated with lower use of insulin the exit time point (E). *p<0.05, ** p<0.01, *** p<0.001

We also questioned whether a patient’s baseline age ratio was correlated with the use of insulin use by their exit from the study. Interestingly, we found that a higher age ratio at baseline, correlated with less insulin use (**Figure 7C, E, Source 6**) and this was not due to differences in CA (**Figure 7B, D, Source 6**).

Overall, we show that pancreatic islet and whole blood tissues have accelerated cellular aging in individuals with T2D with no changes detected in adipose or skeletal muscle revealing the disease affects tissues differently. Cardiovascular, glycemic, and lipid biomarkers, along with adiposity, lifestyle, individual characteristic were found to significantly associate with the rate of aging. Importantly, pharmacological treatment of diabetes, hypertension and hyperlipidemia significantly decreased BA and brought it down to or below CA levels therefore restoring tissue function. We propose that the main driver of accelerated aging in diabetes is the dysregulation in glucose metabolism and low-grade inflammation. While all glycemic, blood pressure, and lipid treatment regimens were found to decrease the rate of aging over 48 months in the ACCORD cohort, intensive regimens reduced the rate of aging to a greater magnitude. No one medication was superior in their ability to reduce the rate of aging over 48 months, and baseline age ratio was not correlated to a higher use of insulin at exit from the ACCORD trial.

## Discussion

Our previous study found that T2D correlated with increased BA at a population level using clinical biomarkers.^4^ and herein this was confirmed using DNA methylation clocks which are thought to be the gold standard to calculate BA. DNAm PhenoAge revealed increased BA in in whole blood and pancreatic islet methylation samples from people with T2D when compared to those without. Gene methylation differences seen between pancreatic islets and whole blood reveal some similarities although the methylation pattern amongst tissues revealed differences that should be further studied to understand the underlying mechanisms of these observations.

Pathway enrichment analysis in pancreatic islets revealed associations with MGMT signaling in DNA damage repair, and many pathways associated with apoptosis, including PAK-2p34, NADE, and SMAC(DIABLO) signaling. Other upregulated pathways included MET, Nectin/Necl, Nodal, and TNFR1 signaling, and hyaluronan biosynthesis. MET signaling inhibits apoptosis and upregulates proliferation and migration.^13^ Upregulation of Nectin/Necl also plays a role in inhibiting apoptosis.^14^ Nodal signaling is involved in the inhibition of beta cell proliferation^15^ and decreases glucose stimulated insulin release. Insulin inhibits nodal signaling and attenuates apoptosis.^16^ TNFR1 mediated ceramide production is involved in impairment of insulin secretion, and apoptosis via ER stress and mitochondrial dysfunction.^17^ Hyaluronan biosynthesis is involved in islet dysfunction, inflammation, and amyloid deposition.^18^

Pathway enrichment in whole blood showed pathways involved in biogenesis of mitochondria, and fibronectin matrix formation. NODAL signaling was also upregulated in whole blood and is involved in maintaining the human pluripotent stem cell population^19^. Upregulation of pathways involved in inflammation were also shown, such as the AIM2 inflammasome^20^, which is involved in inflammatory signaling and inflammatory related apoptosis.

Many of these pathways are relevant to the fundamental biology of tissue aging, such as senescence, mitochondrial dysfunction and stem cell exhaustion which is an initial insight as to which molecular pathways underlie the associations of increased BA in T2D. Further experimental confirmation of these is required.

Surprisingly, there were no differences in DNAm in skeletal muscle and adipose tissue which could be due to their uniquely greater capacity for metabolism of glucose. Adipose and skeletal muscle tissues express GLUT 4 receptors in response to insulin secretion. The upregulation of GLUT 4 receptors increases the uptake of glucose allowing for metabolism or storage in the tissue. It may be that their improved ability to handle large amounts of glucose contribute to decreased damage, or glucotoxicity, decreasing the susceptibility to tissue injury and aging. The accelerated aging in pancreatic islets agrees with the well-described glucotoxicity mechanisms that occur in the setting of T2D.^21^ It has also been shown that insulin resistance plays a role in the senescence of pancreatic β-cells.^22^ Increased aging in whole blood tissues may be due to low grade inflammation in T2D, which is evidenced by the correlation of increased segmented neutrophils, globulin and platelets to groups of faster agers (**Figure 4, Source 4**) along with previous studies noting the inflammatory processes in T2D.^23^

In the paper describing the development of DNAm PhenoAge (Levine et. al. 2021), DNAm PhenoAge had a high correlation with all tissues (0.71). Adipose (0.4) and skeletal muscle (0.52) had a lower correlation with DNAm Phenotypic age, while white blood cells (0.92) had a strong correlation. The pancreas was not included in their analysis. The lower correlation with adipose and skeletal muscle may have also been a cause of the absence of significance in DNAm Phenotypic Age between nondiabetic and T2D individuals.

Herein, we found that glycemic indices had a strong correlation with BA. Our previous data showed an acceleration in aging in both T1D and T2D. Our current data shows that as glycohemoglobin and fasting glucose were increased, the speed of aging was increased. We also saw that the older the individual was when diagnosed with diabetes, the slower their speed of aging was (**Figure 2, Source 2**). This supports our hypothesis that the dysfunction in metabolism of glucose may be the leading driver in accelerated aging in those with diabetes, irrespective of the pathophysiology of disease, which is different in T1D and T2D. The longer an individual has diabetes likely correlates with greater susceptibility of organs to damage by hyperglycemia, leading to accelerated aging.

Abnormalities in cardiovascular measures were found to correlate with accelerated aging, which has been supported by the literature.^24^ The increase in SBP, DBP, pulse and albumin were all correlated with higher speed of aging within the diabetic population. We speculate that this alteration in cardiovascular measures is primarily the result of microvascular compromise due to age advanced glycation end products (AGEs).^25^ Longer periods of hyperglycemia increase the proportion of AGEs, which may cause damage on the microvasculature, resulting less compliance, or more stiffness, that leads to increases in blood pressure. This will require further experimental testing.

Inflammation has been shown to be associated with T2D.^23^ In this study we show that increases in globulin, segmented neutrophils, and platelet counts are correlated with a faster speed of aging within diabetes. Immunoglobulin is a type of globulin molecule synthesized in the liver and used in the innate and adaptive immune responses. Segmented neutrophils are primary mediators in acute innate inflammation, and platelets are activated by circulating IL-6 in the setting of inflammation.^26^ Given the known role of inflammation in tissue aging due to altered cellular communication, this is an additional mechanism that should be considered when assessing BA in T2D.

Lifestyle factors and population characteristics were also correlated with alterations in aging. It is not surprising that aerobic activity was correlated with decreased BA, as it has been largely associated with protective effects on the body, including slowing vascular aging^27^ and cognitive function^28^ in the elderly. Because vigorous exercise is more closely associated to increases in muscle mass, it is possible that the greater glucose requirements for proper function pulls glucose from the blood to be metabolized in the skeletal muscle, therefore allowing for lower blood glucose and less risk of harmful effects on the rest of the body.

We also found that Hispanics had a slower deceleration in their rate of aging over the 48 month period of treatment in the ACCORD trial than whites. The ACCORD trial enrolled individuals from 77 different locations within the United States and Canada. Because these individuals were only chosen based on their disease status, socioeconomic factors, such as food insecurity, and genetic factors could have played a role in the health maintenance of these groups. Studies have also shown that the average glycohemoglobin level in nondiabetic Hispanic individuals is significantly higher than levels in non-Hispanic whites.^29^ Further analysis should be done to study the cause of this striking difference, and to determine whether alternate measures should be considered when diagnosing the Hispanic population with diabetes.

Overall, the female biological sex had a significantly higher dAge at both baseline and 48 months. The biological sex difference in biological aging in people with T2D is interesting since women tend to have a longer lifespan even though they are frailer and have worse health at the end of life.^30^ Whereas the biological mechanisms are not entirely clear, a recent review points to the following factors playing an important role in the sex specificity in human biological aging: sex chromosomes, hormones, mitochondria linked mechanisms such as senescence, telomeres, epigenetics among others.

Interestingly, we found that although many patients had clinical biomarker values within the normal clinic ranges, there were large differences between individual speeds of aging even within those ranges (Figures 2, 3, and 4, Source 2, 3, and 4). While these values would not warrant further follow up within the clinic, we show that patients can have great differences in their BA, and therefore, we suspect differences in their mortality and morbidity. This underscores how BA could be used in the clinic as a more precise measurement of mortality and morbidity than individual clinical biomarkers.

Given the higher burden that T2D has in underrepresented populations, the potential genetic and sociodemographic factors behind the increased BA in Blacks and Hispanics with T2D even after pharmacological treatment need to be studied further such that adequate and specific interventions can be developed to lessen the disease burden.

A key question addressed in this study is whether BA and therefore mortality risk can be altered with treatment. Age ratio within the population as a whole was greater than 1.6 in all subsets of cohorts. After 48 months of glycemia and lipids or blood pressure treatment, age ratio was reduced in all patients. Some individuals had a BA that matched their CA after this time, and many had a BA that became less than their CA. Overall, intensive blood pressure and glycemia cohorts were shown to have a BA that was significantly lower than the BA of individuals in the standard treatment arms. Lipid fibrate, those treated with a fibrate and statin, were shown to have a significantly lower BA than those on the lipid placebo trial, those treated with only a statin. The lipid fibrate cohort had the most reduction in BA over 48 months, and intensive BP trails followed. This data provides support for the legacy effect, also known as metabolic memory, which describes long term positive health outcomes as a result of glycemic control. The underlying mechanisms for this effect continue to be in discussion. BA in these patients who underwent more intensive treatment had almost complete reversal of their BA to match their CA, or in some cases had an even lower BA. The improved microvascular outcomes seen in those with more aggressive treatment may contribute to decelerated aging and lower mortality and morbidity.^31^ HDL has been shown to have anti-inflammatory properties which are beneficial in the reduction of cardiovascular risk, which may have contributed to the lipid fibrate trial group greater deceleration of aging.^32^ However, this seems to be contradicted by our results from the NHANES cohort showing increased aging in those with greater HDL levels and further studies should address this question.

Because mediations have different profiles of side effects on organ systems, we questioned the effect on medications on the rate of aging. For instance, thiazolidinediones have a great side effect profile including, adiposity, osteoporosis, and increased risk of heart failure. All medications reduced age ratio from baseline to exit, and there were no differences in their rates when adjusting for significance that was present at baseline. In both intensive and standard cohorts, greater baseline age ratio was found to correlate to less use of insulin at exit. This was the opposite of what was expected since insulin treatment is usually begun as a result of complete failure of β-cells in T2D.

The differences in aging between tissues should first, be completed at a larger scale and in a greater variety of tissue types, and second, should be compared between individuals to determine if there are differences in the rate of aging between tissues between individuals. This idea could be translated into the clinic as a method of precision medicine and can be considered by physicians when prescribing medication and when following biomarkers of health.

### Study Limitations

Our study had only a limited number of tissue samples used for methylation analysis. In future studies we will repeat this analysis with a greater variety of tissues, especially those affected in diabetes, to show the rates of aging each organ in T2D. Those in the NHANES study who were diabetic were not able to be categorized as a T1D or T2D population. These individuals were pooled as one diabetic population. In the ACCORD Trial, individuals were placed on a treatment regimen that included many different types of medications, therefore making it more difficult to isolate each medication to see its sole effect on aging in T2D. An additional limitation is the lack of mechanistic insight into how BA, either calculated through KDM or DNA methylation, lead to accelerated cellular aging. Whereas this is out of the scope of this work, we are establishing how aging is accelerated in T2D and this can be modified through treatment. Future studies will address the mechanisms that underlie these correlations.

In conclusion, DNAm of blood and islets from people with T2D age faster than those without. Cardiovascular, renal, liver, and lifestyle and individual characteristics correlate with the speed of aging, and more intensive treatment protocols correlate with greater decelerations in the rate of aging. This work contributes to laying the foundation of T2D as a disease of accelerated cellular aging leading to novel pathophysiological pathways and treatment targets.

## Acknowledgements and Funding

This study was supported by Institutional Startup Funds to C.A.M. (Joslin Diabetes Center) and NIH grants P30 DK036836 Joslin Diabetes Research Center (Bioinformatic Core).

## Supplemental Figures

**Supplemental Figure 1, Source 3:**
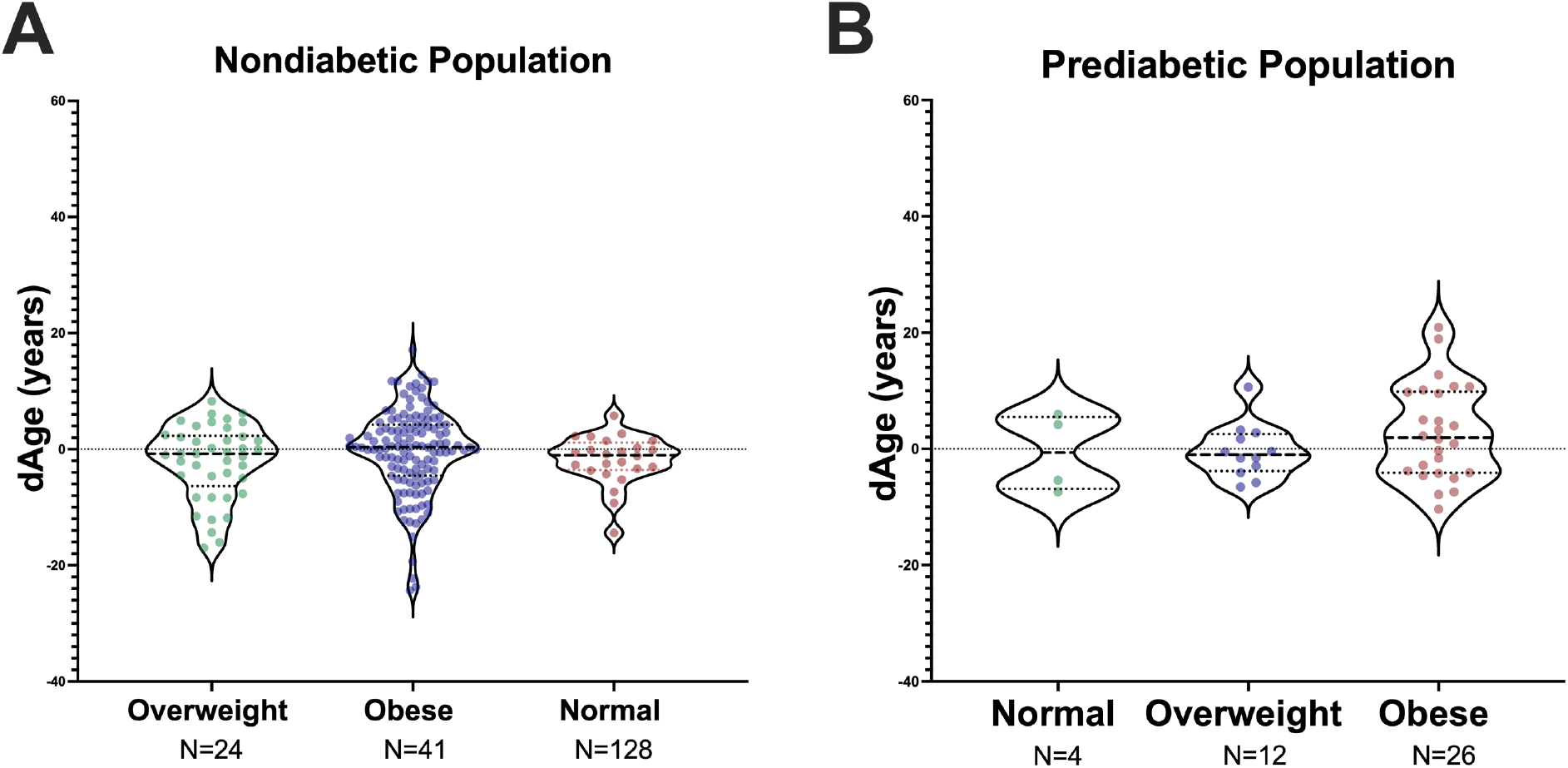
No association between dAge and obesity status in individuals without diabetes (A) and prediabetes (B).

**Supplemental Figure 2.**
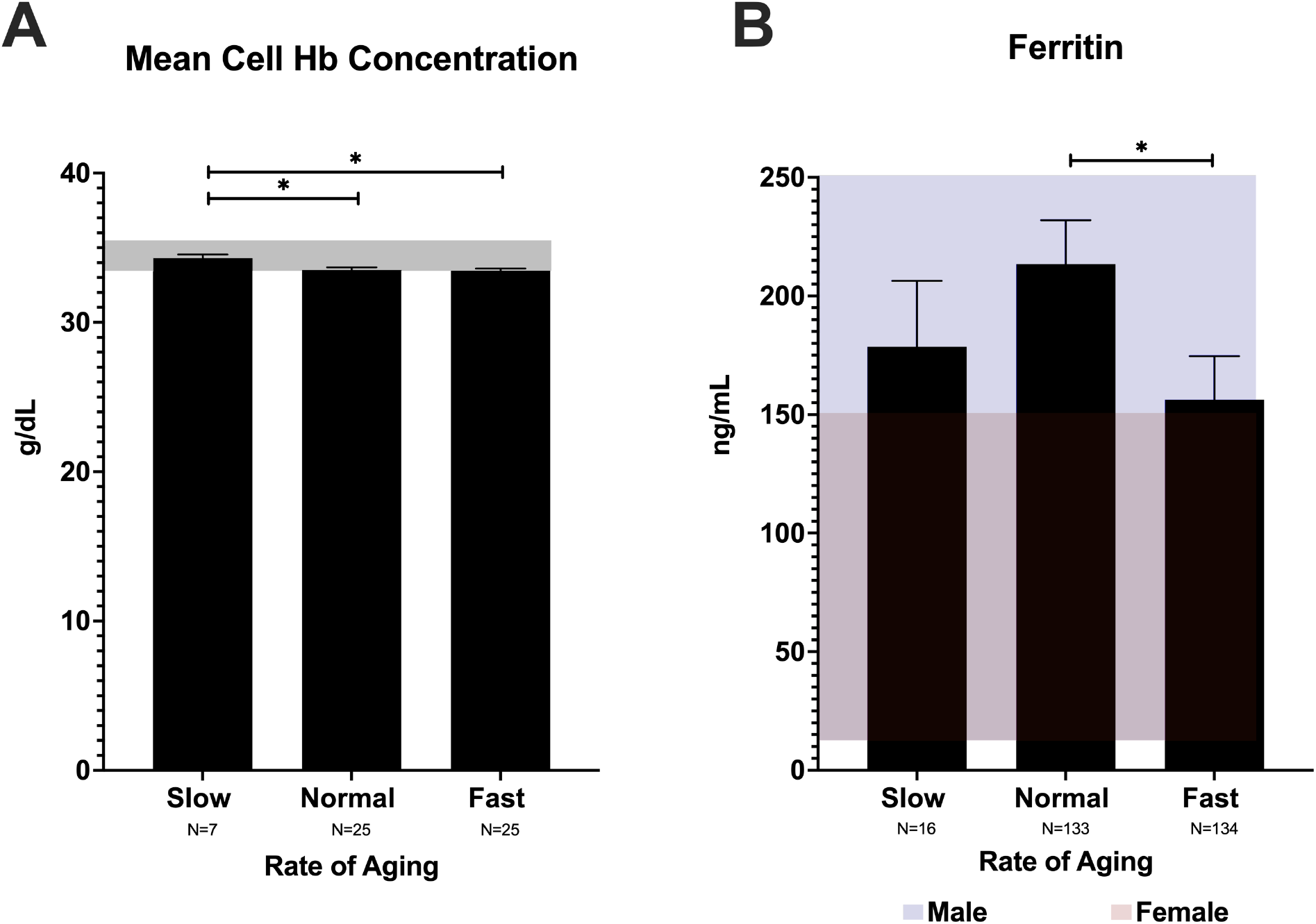
Source 1s: Decreased mean cell hemoglobin (Hb) concentration correlates with faster aging (A). Lower levels of ferritin correlate with faster aging when compared to normal agers (B). *p<0.05, ** p<0.01, *** p<0.001

**Supplemental Figure 3, Source 5:**
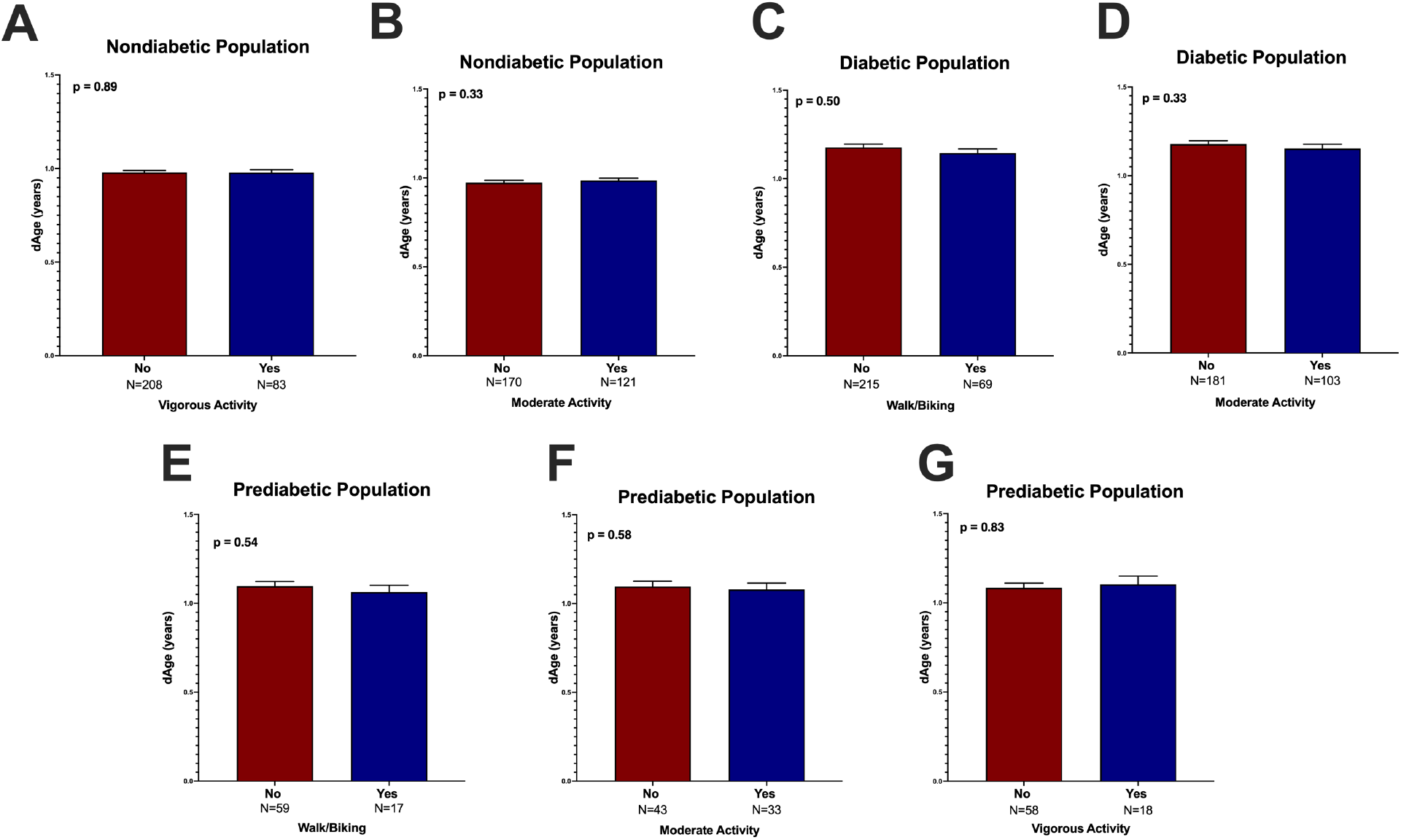
Vigorous activity (A) and moderate activity (B) exercise do not correlate with dAge in those without diabetes. Walking/biking (C) and moderate activity (D) do not correlate with dAge in those with diabetes. Walking/biking (E), moderate (F) and vigorous (G) activity do not correlate with dAge in those with prediabetes.

**Supplemental Figure 4.**
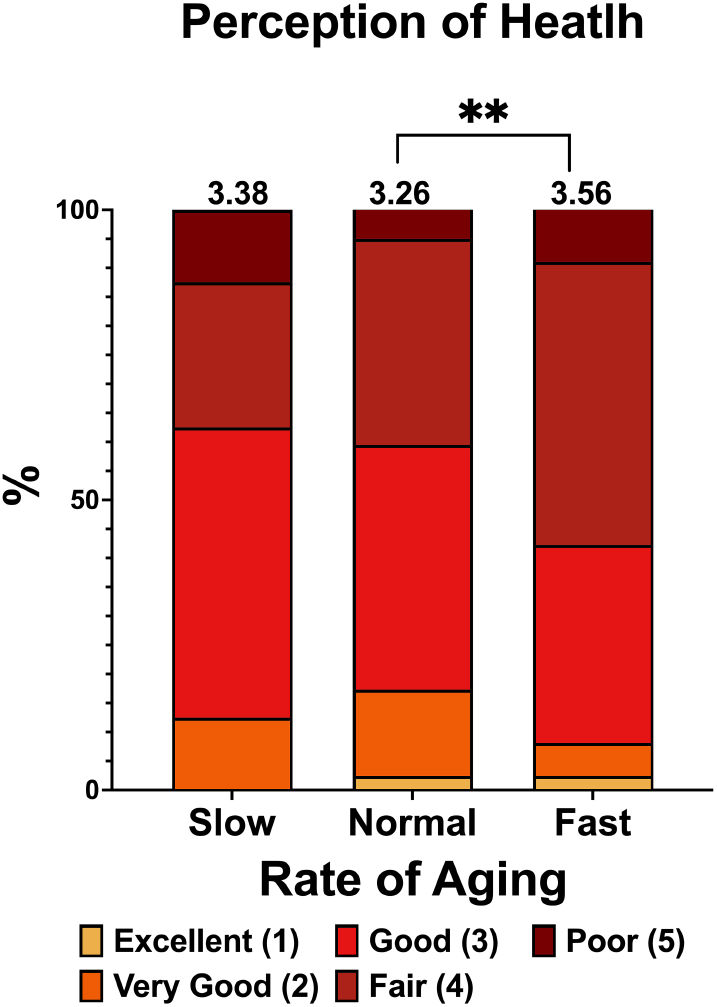
Source 1s: Lower perception of health is correlated with faster rates of aging in the NHANES Diabetic population.

**Supplemental Figure 5, Source 6:**
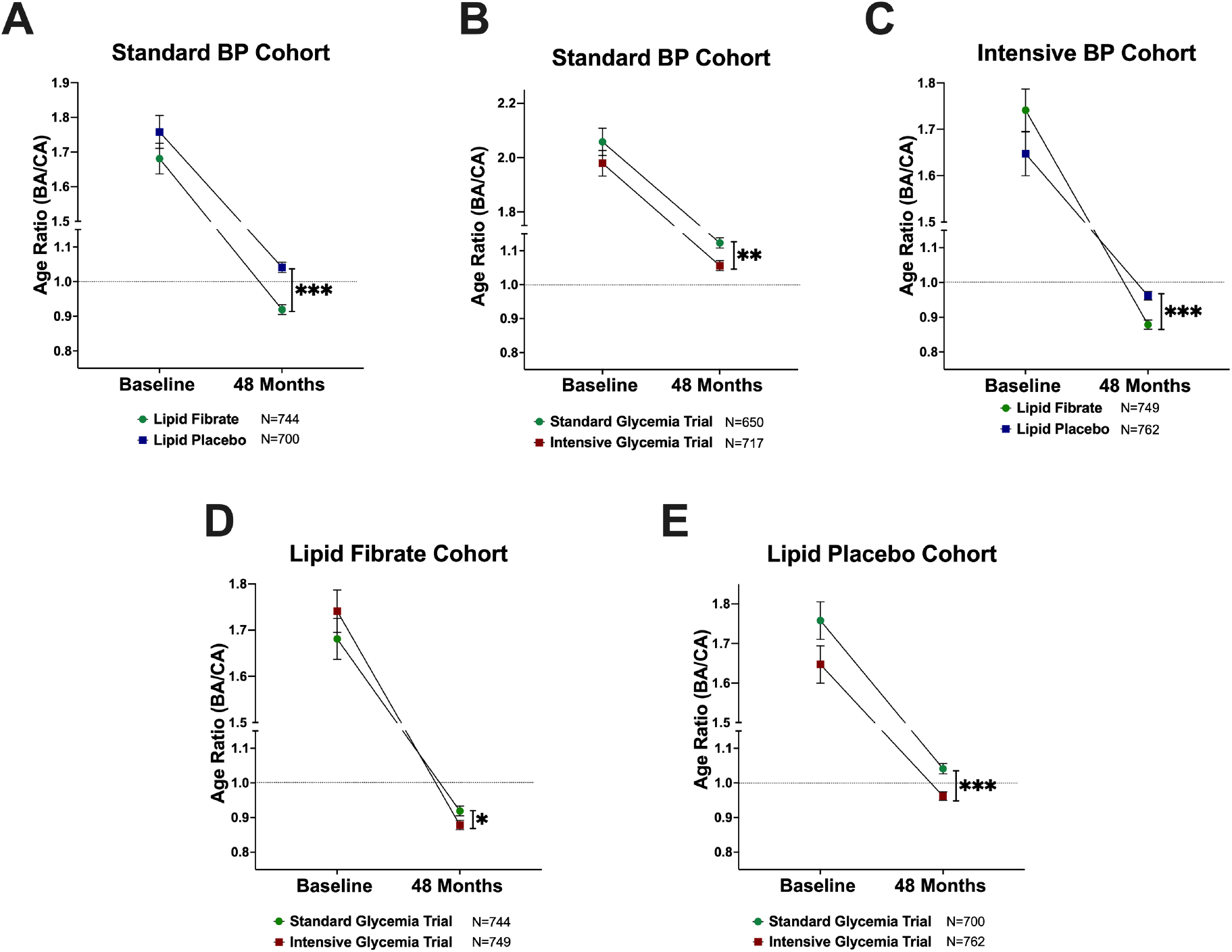
Rate of aging comparisons between ACCORD cohorts after 48 months in the standard blood pressure (A), intensive blood pressure (B and C), lipid fibrate (D), and lipid placebo (E) cohorts. Analyzed using nonparametric t-test. *p<0.05, ** p<0.01, *** p<0.001

